# Distinct RNA engagements define genome import and replication elongation in alphaviruses

**DOI:** 10.64898/2026.07.03.736289

**Authors:** Yaw Bia Tan, Yongbo Luo, Ping Liu, Andres Mertis, Dahai Luo

## Abstract

Membrane-associated replication complexes (RCs) are a hallmark of alphaviruses, yet the mechanisms by which viral RNA is delivered into these compartments and coordinated with RNA synthesis remain incompletely understood. Here, we present a cryo–electron microscopy (cryo-EM) structure of the Chikungunya virus (CHIKV) RC core, consisting of nsP1, nsP2, and nsP4, in complex with the replicative RNA substrates, revealing two spatially distinct RNA-binding sites. A single-stranded RNA (ssRNA) engages the nsP2 helicase and extends toward a pore formed at the nsP1–nsP4 interface, whereas a double-stranded RNA (dsRNA) is accommodated within the central catalytic pocket of the nsP4 polymerase in an elongation configuration. Structural analysis suggests a helicase-assisted RNA threading model in which the 5′ end of the viral genome is guided toward the spherule lumen through the RC pore. Concurrently, the nsP4 polymerase engages dsRNA in a manner consistent with RNA synthesis. This work provides a structural framework for understanding RNA trafficking and enzymatic coordination in viral replication organelles.

**Significance Statement:** Like many positive-sense RNA viruses, alphavirus replication occurs within membrane-bound spherules, but how viral RNA is delivered into these compartments and coupled to RNA synthesis has remained unclear. Here, we show that the CHIKV RC core contains two functionally distinct RNA-engagement modules: an nsP2-associated ssRNA path directed toward the nsP1–nsP4 pore, and an nsP4-bound duplex RNA positioned in an elongation-like polymerase channel. These observations support a model in which alphavirus RCs integrate genome import and RNA synthesis within a single membrane-associated molecular machine. This work provides a structural framework for understanding RNA trafficking, replication-organelle function, and potential antiviral targeting of alphavirus replication.

## Introduction

Of the Alphavirus genus in the Togaviridae family, CHIKV is re-emerging as a clinically important mosquito-borne virus affecting millions of people in subtropical and tropical regions. CHIKV infection may cause acute febrile illness, whereas chronic infection can lead to polyarthritis persisting for years after infection [1-3]. Although vaccines have recently been approved, including the live-attenuated IXCHIQ and the virus-like particle–based VIMKUNYA [4-7], there are currently no specific antiviral therapies. Progress in antiviral development targeting alphavirus enzymes has been stalled by the limited knowledge of the molecular organization of membrane-associated multicomponent alphavirus replication complex (RC), which orchestrates the time-sensitive viral replication process.

Replication organelles (ROs) composed of viral and host materials are common among positive-strand RNA viruses, including coronaviruses, nodaviruses, and alphaviruses [8-12]. These ultrastructures are viral RNA replication centers that localize to virus-induced membrane-associated compartments packed with internal viral RNA condensate and can be located in different subcellular locations, like at the plasma membrane (PM), mitochondrial outer membrane (MT) and endoplasmic reticulum (ER). However, the biogenesis and exact molecular mechanisms that drive replication are not fully understood. For example, little is known about events such as early alphavirus RC biogenesis involving nonstructural proteins (nsPs) autoprocessing, host factor recruitment, RNA import and export at the ROs, initiation of viral de novo RNA synthesis, switching between replication and transcription mode and coordination of enzymatic activities.

The positive-strand RNA (+RNA) genome of CHIKV is approximately 11.8 kilobases long and contains a 5’-cap-0 structure. It resembles a messenger RNA (mRNA), comprising 5’ and 3’ untranslated regions (UTRs), an intergenic region and a polyadenylated 3’ end. The first open reading frame (ORF) of CHIKV genome encodes four nonstructural proteins (nsP1-4) that play enzymatic and non-enzymatic roles. nsP1 is the capping enzyme with methyltransferase and guanylyltransferase activities required for modification of the 5’-end of newly synthesized viral positive strand RNAs. The largest protein nsP2 consists of an N-terminal RNA helicase domain and C-terminal protease domain. nsP3 is known for ADP-ribosylation activity in its macro domain, and nsP4 consists of alphavirus-unique N-terminal domain (NTD) and an RNA-dependent RNA polymerase (RdRP) domain responsible for de novo viral RNA synthesis and polyadenylation [8, 13-15].

Following translation, the nsPs are first presented as polyprotein precursors P1234 and P123, which are subsequently autoprocessed by the nsP2 protease. This process is required to regulate biogenesis of an alphavirus RC embedded in membrane-compartmentalized ultrastructures called spherules and the synthesis of viral negative and positive strand RNAs. The early RC which likely consists of P123 and nsP4 has been proposed to produce negative-strand RNA (-RNA) which later serves as template for the mature RC, which consists of fully processed individual nsP1-4 subunits to produce positive-strand RNAs[14, 16]. This results in both viral +RNA and -RNA species occupying the same spherule space, likely in the form of double-stranded RNA (dsRNA) intermediate, shielded away from host immune sensors.

The structure of the alphavirus RC core was previously determined. In this structure, nsP4 faces the spherule lumen, whereas nsP2 is oriented toward the cytosolic side [8, 17, 18]. However, this structure does not provide mechanistic insight into viral RNA trafficking or replication. Here, we present a cryo-EM structure that captures the nsP2 helicase and nsP4 polymerase bound to a partially double-stranded RNA substrate. This viral replicase RNP structure provides molecular insight into two possible events at the CHIKV replication site: nsP2 helicase–facilitated viral RNA import into the spherule and formation of a viral RNA elongation complex at the nsP4 polymerase. Together, these findings suggest how viral RNA may be handled within alphavirus replication organelles and provide insight into the coordination of enzymatic activities during RNA replication.

## Results

### The RNA-bound alphavirus RC core reveals two spatially distinct RNA-engagement modes

Earlier, we reported a ternary complex of the nsP2 helicase from Chikungunya virus in association with single-stranded RNA, captured in an ATP analog–bound closed conformation [15]. Additional cryo-EM density was detected at the putative RNA pore formed at the interface between nsP1 chain A, nsP1 chain B, and nsP4, a space sufficiently large to accommodate RNA (PDB ID: 7Y38) [8].

To investigate the relevance of these observations to the virus RNA replication process, we modified the in vitro reconstitution of the alphavirus RC core by supplying a high concentration of a replicative RNA, containing a 10-base-pair dsRNA stem–loop region and a 15-nucleotide ssRNA 5′ overhang, throughout the steps of the assembly process (see Methods). This approach enabled us to capture an RNA bound viral RC core structure and reveal two distinct RNA-binding locations (Fig. 1A–D).

**Figure 1.**
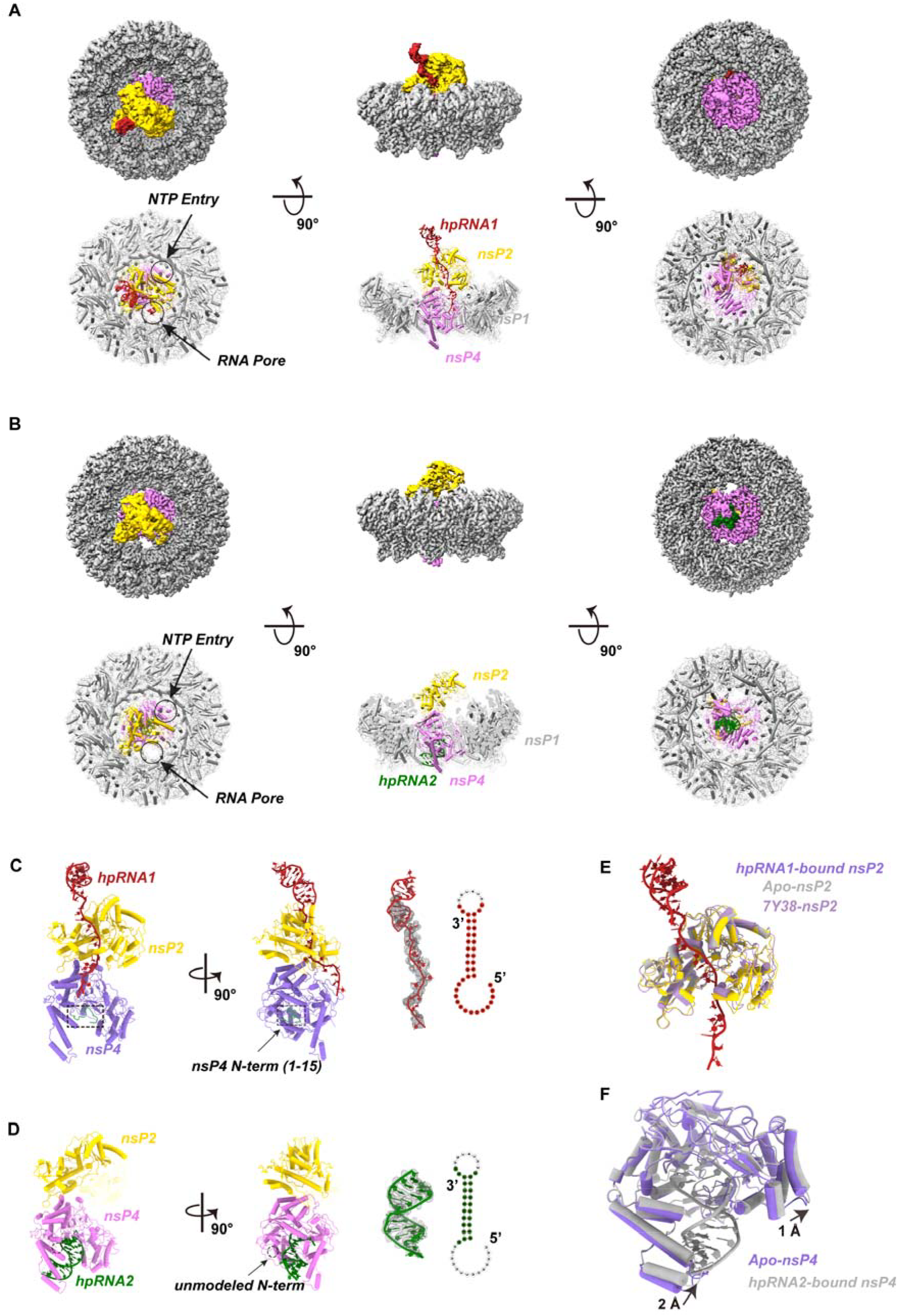
Alphavirus RC core assembly involving two distinct RNA binding events. (A–B) Cryo-EM structure of the alphavirus RC core bound to RNA. The complexes are shown as the cryo-EM density maps in the top rows of (A) and (B), and as atomic models with transparent surfaces in the bottom rows. For both panels, the views are arranged from top, side, and bottom orientations from left to right. (C–D) Two distinct RNA binding events occurring within the alphavirus RC core: (C) binding of hpRNA1 across the nsP2 helicase, and (D) binding of hpRNA2 within the central RNA-binding channel of the nsP4 RdRP. (E) Superimposition of the hpRNA1-bound nsP2 (purple), apo-nsP2 (yellow), and a reference structure (grey) showing allosteric shifts. (F) Subdomain deflections of 2 Å and 1 Å (indicated by arrows) between apo-nsP4 (purple) and hpRNA2-bound nsP4 (grey).

To avoid ambiguity, we refer to the RNA density associated with nsP2 as hpRNA1 and the RNA density bound in the nsP4 polymerase channel as hpRNA2. Both densities arise from particles assembled in the presence of the same designed hairpin RNA substrate. In the reconstructed RC core, however, the nsP2-associated and nsP4-associated RNA densities are spatially separated and are not resolved as a single continuous RNA molecule. We therefore interpret them as two distinct RNA-engagement modes captured within the same RC architecture.

The first RNA-engagement mode is represented by hpRNA1, the ssRNA region of which binds extensively along the RNA-binding groove of the nsP2 helicase (nsP2h) and extends toward the RNA pore, located opposite the nsP4 NTP entry site (Fig. 1C). The 5′ end of the ssRNA is inserted into the RNA pore, suggesting an nsP2h–facilitated RNA import mechanism in which the RNA enters the spherule lumen with its 5′ end leading (Fig. 1A, C). Comparison with the apo nsp2 structure and the previously reported RNA-bound nsp2 structure futher revealed RNA-induced allosteric shift within the helicase domains (Fig. 1E).

The second RNA-engagement mode is represented by hpRNA2, which occupies the central RNA-binding tunnel of the nsP4 polymerase (Fig. 1B, D). In this density, ribonucleotide residues U1-C25 and G36-A45 correspond to the template and nascent strands, respectively (Fig. 1D). The interaction with the dsRNA region involves the N-terminal domain (NTD) and all three subdomains of the nsP4 RdRP domain (Fingers, Palm, and Thumb), whereas the 5’ overhang is exposed to solvent and not clearly resolved, assembling into a viral RNA elongation complex (EC). Structural comparison with the apo nsp4 revealed modest RNA-induced rearrangements, inducing subdomain movement of approximately 1-2 Å upon RAN binding (Fig. 1F).

Together, these distinct ssRNA and dsRNA binding events observed in the CHIKV RC core likely represent key steps in substrate recognition during the viral RNA replication process.

### A putative elongation complex at the nsP4 RdRP within the RC core

The RNA polymerase component of the alphavirus RC core, nsP4, plays an essential role in viral RNA replication by catalyzing de novo RNA synthesis in the 5′-to-3′ direction to produce both +RNA and −RNA. nsP4 comprises an NTD, including a 14–amino acid N-terminal tip (N-tip), and an RNA-dependent RNA polymerase (RdRP) domain. The RdRP domain adopts a canonical right-hand fold and consists of three subdomains—Fingers (further subdivided into Index, Middle, Ring, and Pinky), Palm, and Thumb (Fig. 2A).

**Figure 2.**
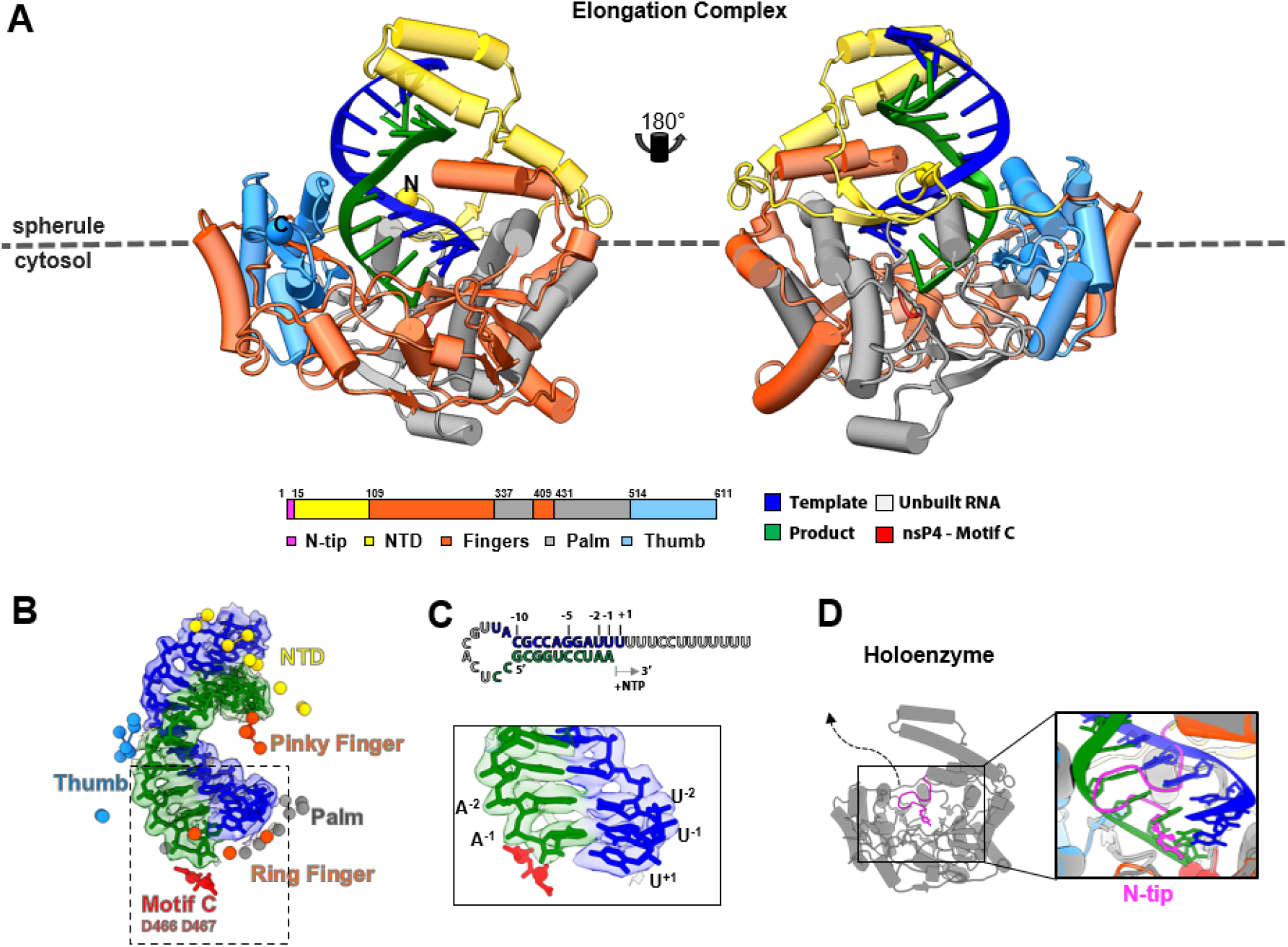
nsP4 binds dsRNA to form a putative elongation complex. (A) RNA-bound nsP4 colored by subdomain: NTD, yellow; Fingers, orange; Palm, gray; and Thumb, light blue. Motif C at the nsP4 active site is highlighted as red C_α_ spheres for residues D466 and D467. The RNA duplex is colored with the template strand in blue and the product strand in green. (B–C) nsP4 residues contacting RNA are shown as C_α_ spheres and colored according to the subdomain scheme in (A). The Fingers subdomain is further subdivided into Index, Middle, Ring, and Pinky fingers; the Ring and Pinky fingers are shown. In (C), the colored RNA sequence (top) corresponds to the dsRNA modeled in the nsP4 channel, and the corresponding cryo-EM density coverage is shown below (bottom). (D) The N-tip region of nsP4 retracts from the RNA-binding channel and the active site upon RNA accommodation.

The assignment of the nsP4-bound duplex RNA as an elongation-like state is based on its position and polarity within the RdRp channel. The duplex RNA occupies the canonical template-product RNA-binding path, with the putative product-strand 3′ end positioned near motif C and the conserved GDD catalytic residues (Fig. 2A–C). This geometry resembles elongation complexes observed in other viral RdRPs. However, we did not observe unambiguous density for an incoming NTP or remdesivir triphosphate at the active site, and the catalytic metal-ion coordination could not be assigned with confidence at the present map resolution. Therefore, we refer to this structure as a putative elongation-like state rather than a catalytically validated elongation complex. The nsP4–RNA interface is extensive (1,539 Å^2^) and involves all three RdRP subdomains. In addition, the alphavirus-specific NTD—harboring a helix–turn–helix (HTH) motif—provides additional stabilization of RNA binding. Notably, the α-helix–rich Pinky finger also contains an HTH-like motif positioned adjacent to the NTD: the Pinky finger contacts the major groove of the dsRNA, whereas the NTD engages the minor groove.

Comparison of the apo and dsRNA-bound nsP4 structures within the RC reveals minimal global conformational changes. The primary difference is localized to the N-terminal tip (N-tip). In the apo structure, the N-tip occupies a position that overlaps with the dsRNA-binding path, whereas it is not resolved in the dsRNA-bound structure, suggesting displacement to accommodate RNA binding (Fig. 2D). These observations suggest that the NTD and Pinky finger of nsP4 may play regulatory roles in RNA binding, translocation, and stabilization during viral RNA replication. Such a mechanism is reminiscent of cofactor-mediated stabilization in the SARS-CoV-2 RdRP complex, where accessory proteins (nsp7 and nsp8) stabilize RNA engagement and enhance processivity [19-21], as well as intramolecular regulatory elements in flaviviral NS5, in which the C-terminal tail and priming elements modulate RNA binding and the transition between initiation and elongation states [22, 23].

### Helicase-guided RNA import via the RNA pore

Helicases are ATP hydrolysis–driven motor proteins that remodel RNA and DNA during essential cellular processes. nsP2 belongs to the SF1 family of viral helicases. It couples the energy derived from nucleotide triphosphate (NTP) hydrolysis to translocate along a single-stranded RNA loading strand, actively unwinding double-stranded RNA (dsRNA) intermediates in the 5’ to 3’ direction [15, 24]. In our structure, the ssRNA portion of the supplied hairpin substrate traces a continuous path from the nsP2 helicase (nsP2h) toward the RNA pore located at the interface between the nsP1 hooking loop (chains A and B) and the dominant region of the nsP4 Fingers subdomain (Fig. 3). ssRNA engagement involves residues from both nsP1 and nsP4 and is accompanied by a modest increase in positive electrostatic potential within the pore upon RNA binding (Fig. 3A–B). The nsP2h N-terminal domain (NTD), which contacts nsP4, appears to constrain nsP2h orientation and thereby impose a 3′-to-5′ translocation polarity that would guide the 5′ end of the RNA into the pore (Fig. 3C–D). Together, our structural data suggest a helicase-facilitated RNA import mechanism in which the viral genomic RNA is threaded into the replication spherule to initiate RNA replication. Such an import process could also correlate with the establishment of the spherule during RC biogenesis [9].

**Figure 3.**
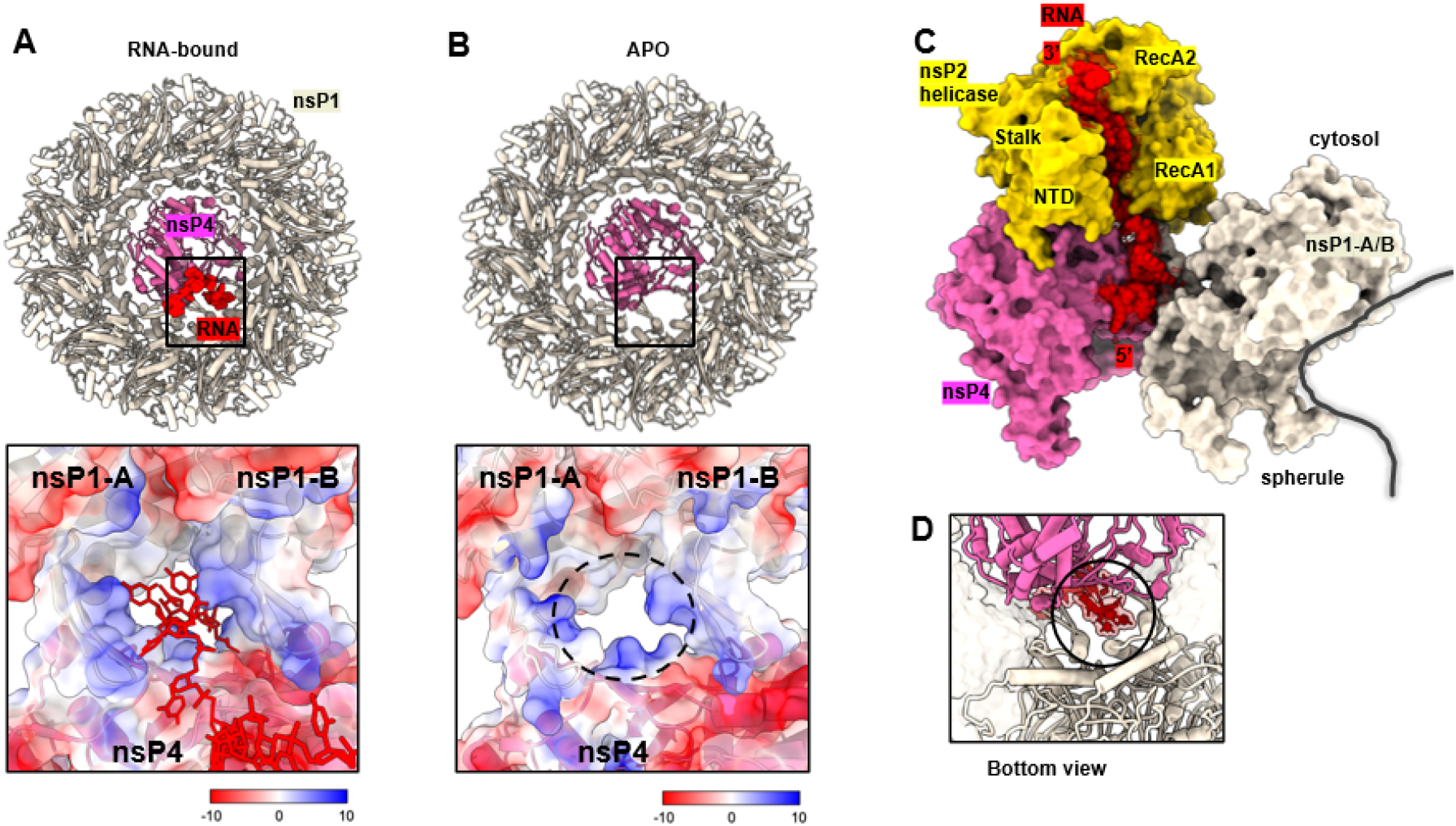
Helicase-guided RNA import through the CHIKV RC pore. (A-B) Bottom views of the RNA pore formed by nsp1 and nsp4 in the RNA-bound (A) and apo (B) RCs. Upper panels show overall structures; lower panels display magnified electrostatic surface representations. (C) Surface representation of the ssRNA-bound nsP2 helicase in relation to the RNA pore formed by nsp1 and nsp4, illustrating a continuous RNA path. (D). Enlarged bottom view highlighting the 5’ overhang of the ssRNA within the pore.

## Discussion

A topological problem is created when positive-strand RNA viruses replicate their genomes inside membrane-associated replication organelles: viral RNA must be selectively recruited, copied, protected from host sensors, and routed for translation or packaging [25-29]. Our findings provide structural insights into understanding how alphaviruses may solve part of this problem. We report two spatially distinct RNA-engagement modes: an ssRNA path along nsP2 directed toward the nsP1–nsP4 pore, and a duplex RNA positioned within the nsP4 polymerase channel in an elongation-like state. Although this structure does not represent a complete continuous replication trajectory, it suggests that the alphavirus RC is not merely a membrane-associated enzyme assembly, but an integrated RNA-trafficking and RNA-synthesis machine.

The molecular organization of the CHIKV RC core differs substantially from other +RNA virus replicases (Fig. 4). Flavivirus NS3 and NS5 form helicase–polymerase assemblies that coordinate RNA synthesis [23], but these complexes are not organized around an nsP1-like membrane-associated pore (Fig. 4C). In coronaviruses, the polymerase–helicase replication-transcription complex and the DMV-spanning RNA translocation pore are structurally and compositionally distinct [30] (Fig. 4D). By contrast, the CHIKV RC core places the nsP2 helicase, nsP4 polymerase, and nsP1-associated pore within a continuous membrane-associated architecture. This compact arrangement may be particularly suited for coupling directional RNA movement at the spherule neck with RNA synthesis inside the ROs [31].

**Figure 4.**
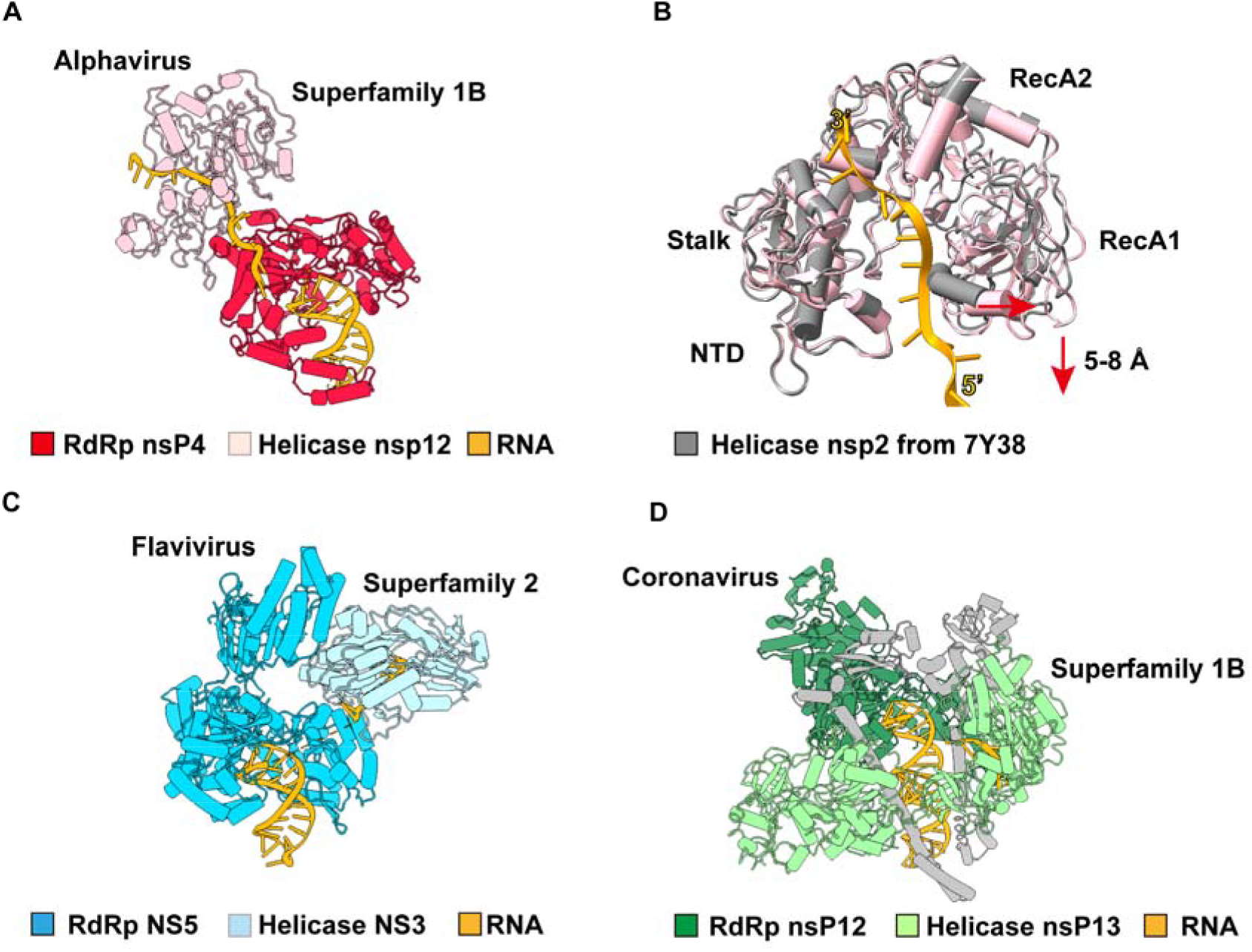
Coordinated organization of helicase and polymerase in CHIKV and other positive-strand RNA virus. (A) Engagement of CHIKV nsP4 polymerase and nsP2 helicase with ssRNA, as observed in this study. (B) Conformational changes in the CHIKV nsp2 helicase upon RNA binding. The previously reported mature RC structure (PDB ID: 7Y38) is shown for comparision. (C) Cryo-EM structure of the Dengue virus NS5-NS3 elongation complex. (PDB ID: 8GZR) (D) Cryo-EM structure of the SARS-CoV-2 replication-transcription complex bound to nsp13 helicase (nsp12-nsp8-nsp7-nsP13) (PDB ID: 6XEZ).

Within this architecture, the orientation of nsP2 relative to the nsP1–nsP4 pore positions the bound ssRNA such that its 5’ end is directed toward the pore. This supports a vectorial RNA-threading model, although the precise relationship between helicase translocation polarity, RNA strand orientation, and pore-directed import will require further biochemical validation. In parallel, nsP4 accommodates duplex RNA within its central polymerase channel, with the nascent strand positioned near the conserved GDD active-site motif, consistent with an elongation-like state. Thus, the structure captures two mechanistically distinct RNA-handling events that may be coordinated during alphavirus replication: helicase-associated RNA delivery and polymerase-mediated RNA synthesis. The structure provides a framework for formulating experimentally testable models of alphavirus RNA handling. The RC core offers a molecular basis for investigating how genome recruitment, RNA threading, and RNA synthesis may be coordinated at the spherule neck. This framework should guide future studies of replication-organelle assembly, host-factor regulation, and antiviral targeting of alphavirus replication.

More broadly, the CHIKV RC structure illustrates how positive-strand RNA viruses have evolved distinct molecular solutions to the shared challenge of coordinating RNA movement, membrane compartmentalization, and genome replication. Coronaviruses encode an exonuclease-based proofreading system, whereas alphaviruses lack a known proofreading exonuclease [32]. The observed nsP2–nsP4 organization therefore may reflect a replication strategy centered on efficient RNA trafficking and synthesis rather than error correction. Whether the nsP2–nsP4 arrangement influences polymerase fidelity, template reuse, or the balance between negative- and positive-strand RNA synthesis remains an important question for future studies.

## Materials and Methods

### Recombinant protein production and *in vitro* reconstitution of RC

Recombinant full-length CHIKV nsP1 fused to a Strep-3×FLAG tag, full-length CHIKV nsP2, and ONNV nsP4 were produced as previously described [8]. Briefly, nsP2 and nsP4 expression constructs in the pSUMO-LIC vector, encoding an N-terminal hexahistidine–SUMO fusion tag, were expressed in Escherichia coli and purified through a multi-step workflow: affinity purification using Ni-NTA beads (Bio Basic), SUMO protease cleavage to remove the N-terminal fusion tag (1 mg protease per 10 mg recombinant protein), ion-exchange chromatography on a Heparin 5 HP column, and size-exclusion chromatography (SEC) on a Superdex 200 16/600 column (Cytiva). After SEC, recombinant nsP2 and nsP4 were flash-frozen and stored in 25 mM HEPES (pH 7.4), 300 mM NaCl, 5% glycerol, and 2 mM DTT until downstream applications.

Membrane-associated CHIKV nsP1 was expressed in transiently transfected Expi293 suspension cultures. The recombinant protein was extracted using buffer A (50 mM HEPES, pH 8, 150 mM NaCl, 5% glycerol, 2.5% sucrose, 1% DDM, 1 mM PMSF), immobilized on Strep-Tactin Sepharose beads (IBA), and washed with assembly buffer B (50 mM HEPES, pH 8, 150 mM NaCl, 5% glycerol, 2.5% sucrose, 0.01% GDN, 2.5 mM magnesium chloride, 1 mM TCEP) prior to binding nsP2 and nsP4 to form the RC core. Following overnight incubation, the assembled RC core complexes were eluted with buffer B supplemented with 10 mM desthiobiotin.

### Cryo-EM sample preparation

Approximately 0.7–1.0 µM RC was mixed with 10–20 µM RNA (5′-UUUUUUUCCUUUUUUUUAGGACCGCAUUGCACUCCGCGGUCCUAA-3′) and 0.5 mM remdesivir triphosphate (RTP) in reaction buffer (50 mM HEPES, pH 8.0, 150 mM NaCl, 1.25% sucrose, 0.01% GDN, 5 mM magnesium acetate, 2.5 mM manganese chloride, 5 mM TCEP) to a total volume of 300 µL and incubated for 2h at ∼25 °C. The mixture was concentrated to 20 µL and supplemented with 5 µL of an RTP : Mg cocktail (to a final concentration of 0.5 mM RTP and 1 mM Mg), followed by a 10 min incubation at ∼25 °C. Five microliters of the final mixture was applied to a glow-discharged Quantifoil R1.2/1.3 300-mesh gold grid coated with monolayer graphene and incubated for 3 min in a Vitrobot chamber equilibrated to 100% relative humidity at 4 °C. The grid was blotted with a −2 blot force for 5 s and plunge-frozen in liquid ethane cooled by liquid nitrogen.

### Cryo-EM data collection and structural determination

Data were collected from the selected grid using EPU on a Titan Krios equipped with a Gatan K2 detector coupled to a BioContinuum energy filter. Detailed acquisition parameters are summarized in Table S1. Movies stacks were preprocessed using Warp or CryoSPARC Patch Motion Correction and Patch CTF Estimation. All downstream processing was performed primarily in CryoSPARC v4.7 [33]. Particles were initially picked using a combination of manual picking and Topaz-based automated picking, followed by duplicate removal, yielding 240,235 particles. After one round of 2D classification, 215,076 particles from well-defined classes were selected for further analysis. These particles were subjected to heterogeneous refinement using six low-pass filtered initial models (20–80 Å) to remove junk particles and resolve structural heterogeneity. The best-resolved class (60.7%) was selected and refined using homogeneous refinement, followed by 3D classification. A subset of 141,832 particles yielded an initial reconstruction at 3.21 Å resolution and was further classified into three clusters. Two major clusters (44.4% and 26.6%) comprising 99,228 particles were combined and refined using homogeneous refinement. To resolve local conformational variability, a focused 3D classification was performed on the nsP2–nsP4 region using a mask specifically targeting this region.

This analysis successfully resolved two major states: Conformation 1 (nsP2-bound hpRNA1; 35,651 particles) and Conformation 2 (nsP4-bound hpRNA2; 35,835 particles). To achieve the final high-resolution reconstructions, Conformation 1 was subjected to an additional round of focused mask creation and 3D classification. The highest-quality sub-class (Class 1; 12,978 particles) was isolated and advanced to global/local CTF refinements, non-uniform refinement, and local refinement, yielding a final density map at 3.42 Å resolution. Conformation 2 was directly subjected to homogeneous, non-uniform, global/local CTF, and local refinements, resulting in a final reconstruction at 3.23 Å resolution. Maps were also post-processed with DeepEMhancer [35] to aid structure interpretation. The final model was built and refined using a combination of manual and automated approaches in Coot [36], Phenix [34], ISOLDE [37], and ModelAngelo [38].

## Data availability

The single-particle analysis cryo-EM density maps of the Chikungunya virus nsP1-nsP2-nsP4 replication complex bound with hpRNA1 and hpRNA2 have been deposited in the Electron Microscopy Data Bank (EMDB) under the accession codes EMD-81199 (Conformation 1) and EMD-58420 (Conformation 2). The corresponding atomic coordinates have been deposited in the Protein Data Bank (PDB) under the accession codes 27JE (https://doi.org/10.2210/pdb27je/pdb) and 31HT (https://doi.org/10.2210/pdb31ht/pdb).

## Author Contributions

Yaw Bia Tan and Dahai Luo designed the study; all authors performed the experiments and analyzed the data; Yaw Bia Tan, Ping Liu, Yongbo Luo and Dahai wrote the manuscript with inputs from all authors.

## Acknowledgments

We thank the members of our laboratories for their helpful discussions and insightful comments on the project. This research was supported by the Singapore Ministry of Education (MOE) under its MOE Academic Research Fund (AcRF) Tier 2 Award (MOE-T2EP30220-0009), Singapore Nucleic Acid Therapeutics Initiative (NATi) N07 grant H25J5a0061, and the National Institute of Allergy and Infectious Diseases of the NIH under award number R01AI187483 to D.L. This work was supported by grants from Estonian Research Council (PRG1154 and PRG3145) to A.M. The funders had no role in study design, data collection and interpretation, or the decision to submit the work for publication. The authors declare no conflict of interest.

