## Supplemental table and figure for "Distinct RNA engagements define genome import and replication elongation in alphaviruses"

*Dahai Luo

**This file includes:**

Figure S1

Table S1

Supplementary References

**Figure S1 Cryo-EM analysis of nsP1+2+4+RNA RC core.** (A) cryo-EM workflow in cryosparc at C1 refinement symmetry. (B) Angular distribution of particles used for the final 3D reconstruction (C) Local resolution distribution of the final map (D) GSFSC curves of local 3D refinement obtained from cryoSPARC.

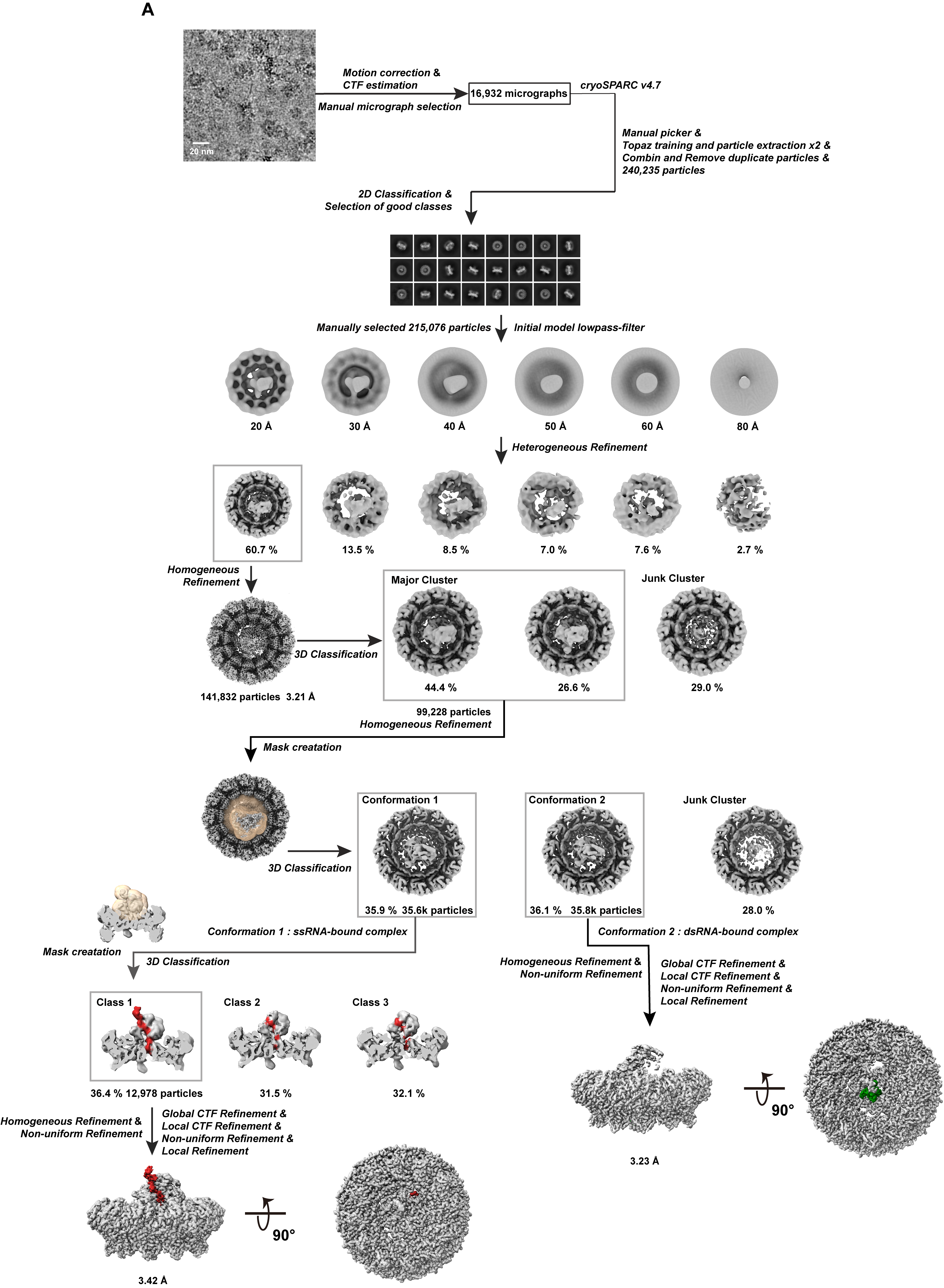

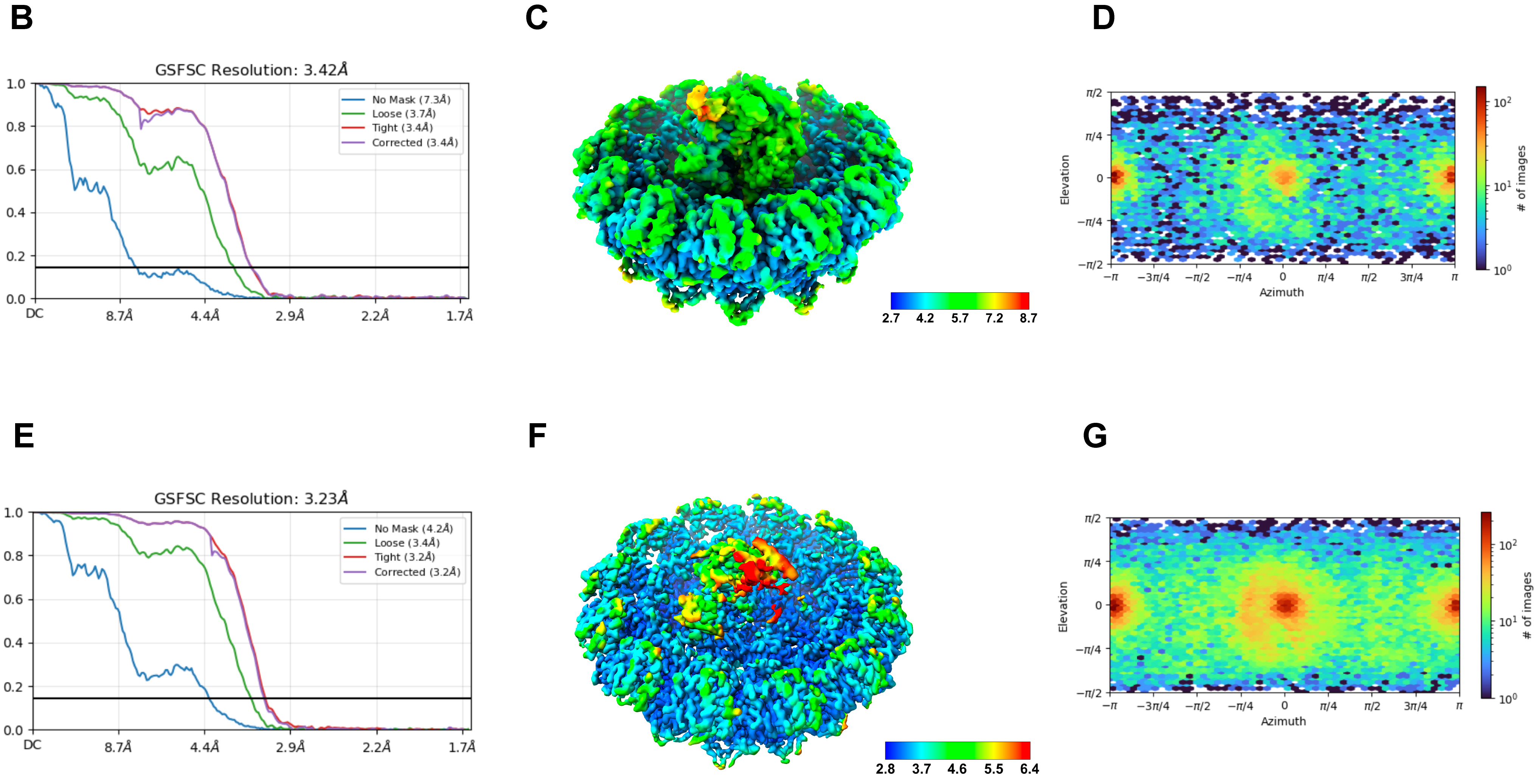

**Table S1**

|  | **nsP1-nsP2-nsP4 replication complex bound with hpRNA1** | **nsP1-nsP2-nsP4 replication complex bound with hpRNA2** |
| --- | --- | --- |
|  | **EMDB: 81199** | **EMDB: 58420** |
|  | **PDB: 27JE** | **PDB: 31HT** |
| **Data collection and processing** |  |  |
| Detector | Gatan K2 | |
| Magnification | 130,000 × | |
| Voltage (kV) | 300 | |
| Electron exposure (e–/Å^2^) | 34 | |
| Defocus range (µm) | -0.5 ~-3.2 | |
| Pixel size (Å) | 1.10 | |
| Symmetry imposed | C1 | |
| Initial particle images (no.) | 240,235 | |
| Final particle images (no.) | 12,978 | 35,835 |
| Map resolution (Å) | 3.42 | 3.23 |
| FSC threshold micrographs | 0.143 | 0.143 |
| Map resolution range (Å) | 2.7-8.7 | 2.8-6.4 |
| **Refinement** |  |  |
| Initial model used (PDB code) |  |  |
| Model resolution (Å) | 2.2 | 2.2 |
| FSC threshold | 0.143-0.5 | 0.143-0.5 |
| Model resolution range (Å) | 1.7~3.3 | 1.6~3.2 |
| Map sharpening B-factor (Å^2^) | 93.3 | 113.2 |
| **Model composition** |  |  |
| Non-hydrogen atoms | 52,127 | 50,585 |
| Protein residues | Protein: 6531 | Protein: 6372 |
| Nucleotide | 39 | 24 |
| Water | 0 | 0 |
| Ligands | ZN: 12 | ZN: 12 |
| B factors (Å^2^) |  |  |
| Protein | 32.49/214.74/77.55 | 35.18/223.12/80.78 |
| Nucleotide | 86.11/318.36/240.55 | 59.48/173.22/96.31 |
| Water | --- | --- |
| Ligand | 108.92/136.43/123.79 | 102.17/131.40/117.4 |
| **R.m.s. deviations** |  |  |
| Bonds length (Å) | 0.003 | 0.003 |
| Bonds Angle (˚) | 0.592 | 0.562 |
| **Validation** |  |  |
| MolProbity score | 2.13 | 1.96 |
| Clashscore | 5.15 | 4.23 |
| Q-score |  |  |
| Rotamer outliers (%) | 3.56 | 2.93 |
| **Ramachandran plot** |  |  |
| Favored (%) | 93.19 | 93.97 |
| Allowed (%) | 6.70 | 5.95 |
| Disallowed (%) | 0.11 | 0.08 |
